# Protein tyrosine phosphatase receptor type D - A convergent risk gene for MDD, BPD, and ADHD and regulator of dopaminergic neuroplasticity in reward associated circuits; A multivariate GWAS, brain tissue and cell type enrichment, and gene fine-mapping

**DOI:** 10.1101/2025.09.17.676918

**Authors:** Christopher Lawrence, Thomas F. Hansen

**Affiliations:** Wake Forest Department of Biology, Winston-Salem, NC, USA; Information Systems and Wake Forest University. WFU High Performance Computing Facility, Winston-Salem, NC, USA; DIS Research, Copenhagen, Denmark; Neurogenomic, Translational Research Centre, Copenhagen University Hospital, Glostrup, Denmark

## Abstract

Major depressive disorder (MDD), bipolar disorder (BPD), and attention-deficit hyperactivity disorder (ADHD), are prevalent, highly heritable psychiatric disorders with significant degrees of genetic overlap. Using open-sourced summary statistics from the Psychiatric Genomics Consortium and 1000 Genomes European reference panel, we utilize a plethora of statistical frameworks to estimate shared genetic liability across MDD, BPD, and ADHD. We observed significant enrichment of latent genetic variables in dopamine neurons, and identify protein tyrosine phosphatase receptor delta (*PTPRD*), as a candidate gene. Literature review and gene fine-mapping suggest that *PTPRD* indirectly regulates *DAT* transporter surface trafficking and neuronal excitability. We propose that the loss of PTPRD’s D1 phosphatase domain results in hyperactivated tropomyosin receptor kinase B (*TrkB*) and rearranged during transfection proto-oncogene tyrosine-protein kinase receptor (*RET*) signaling in dopamine neurons, potentiating excessive downstream phosphorylation and activation of *VAV2*, *ERK1/2*, and *PKC*. We hypothesize that these mechanisms contribute to the significantly altered reward-related behaviors such as cocaine-conditioned place preference, cocaine self-administration, motivation for cocaine, and goal oriented behavior that are consistently observed across *PTPRD* KO/inhibition mice studies. With the inclusion of literature review, we present the first integrated multivariate genetic and mechanistic analysis aimed to identify pleiotropic, cell type-specific mechanisms driving psychiatric comorbidity at the biochemical level.

## Introduction

The World Health Organization estimates that 5% of adults suffer from major depressive disorder globally (World Health Organization, 2023). The Diagnostic and Statistical Manual of Mental Disorders, 5th edition, characterizes major depressive disorder as having 5 or more of the following symptoms: depressed mood, loss of interest in pleasurable activities, weight loss or gain, sleep disturbances, psychomotor agitation or retardation, fatigue, feelings of worthlessness or excessive guilt, decreased concentration, and thoughts of suicide or death (American Psychiatric Association, 2025). The etiology of Major depressive disorder is understood to arise from the complex interplay of biological, genetic, environmental, and psychological factors (Bains & Abdijadid, 2023). Genetic factors are strongly implicated in the development of major depressive disorder, and has been thoroughly demonstrated in twin, adoption, and family studies. The latest twin studies estimate the heritability to be between 40 and 50%, and family studies show that first degree relatives have a twofold to threefold higher lifetime risk of developing major depressive disorder (Lohoff, 2010).

The global prevalence of ADHD in adults and children is estimated to be 5% (The diagnosis and management of ADHD (Bowling & Nettleton, n.d.). According to the Diagnostic and Statistical Manual of Mental Disorders, 5th edition, ADHD is diagnosed on the basis of the amount and severity of symptoms associated with inattention, hyperactivity, and impulsivity. Genetic factors are heavily implicated in the development of ADHD, with a formal heritability of about 80% (Grimm & Kranz & Reif 2020). The latest twin studies have reported heritabilities between 77-88% (Faraone & Larsson).

It is estimated that 2.4% of the global population is affected by bipolar disorder (Zhong et al., 2024). Bipolar disorder is characterized by chronic alternation between manic and major depressive episodes (Nierenberg et al., 2023). Mania is distinguished by periods of unusually high energy and markedly elevated moods or emotions that diverge significantly from one’s normal state. Genetic factors are highly implicated in BPD, with the latest twin studies estimating the heritability to be 79-93% (Barnett $ Smoller, 2013).

Mental disorder comorbidity is an extremely common phenomenon, and lifetime occurrences of mental disorders are highly correlated with increased risk for the onset and development of subsequent mental disorder(s) (McGrath et al. 2020). ADHD especially is a potentiator of comorbidity and is associated with increased MDD and BPD hazard ratios (Kessler et al. 2010). MDD is notably heterogenic, and frequently encompasses characteristic symptoms of bipolar disorder such as mania (Angst et al. 2010). There is extensive genetic overlap across MDD, BPD, and ADHD (Hindley et al. 2023).

GWASs have identified a substantial amount of reproducible risk loci associated with BPD, ADHD, and MDD (Major Depressive Disorder Working Group of the Psychiatric Genomics Consortium, 2025) (Demontis et al. 2023) (O’Connell et al. 2025). Concurrently, there is significant degree of genetic overlap across the three disorders, complicating accurate diagnosis and treatment (Davis et al. 2024) (Donovan & Owen 2016). In this study, we investigate the shared genetic architecture of MDD, BPD, and ADHD, with the aim to examine pleiotropic mechanisms that underlie the development of these psychiatric disorders.

## Methods

The univariate GWAS summary statistics were acquired from the Psychiatric Genomics Consortium (Major Depressive Disorder Working Group of the Psychiatric Genomics Consortium, 2025) (Demontis et al. 2023) (O’Connell et al. 2025). The PGC provides a plethora of open-sourced summary statistics from GWAS, excluding 23andme cohorts. The most recent BPD, ADHD, and MDD GWAS cohorts of European ancestry were used to carry out this study. Practicing strict quality control, we removed all SNPs below a minor allele frequency threshold of 0.01 during summary statistics harmonization. The 1000 Genomes Project Phase 3 panel was used for reference of European genetic variation (Auton et al., 2015).

**Fig 1.**
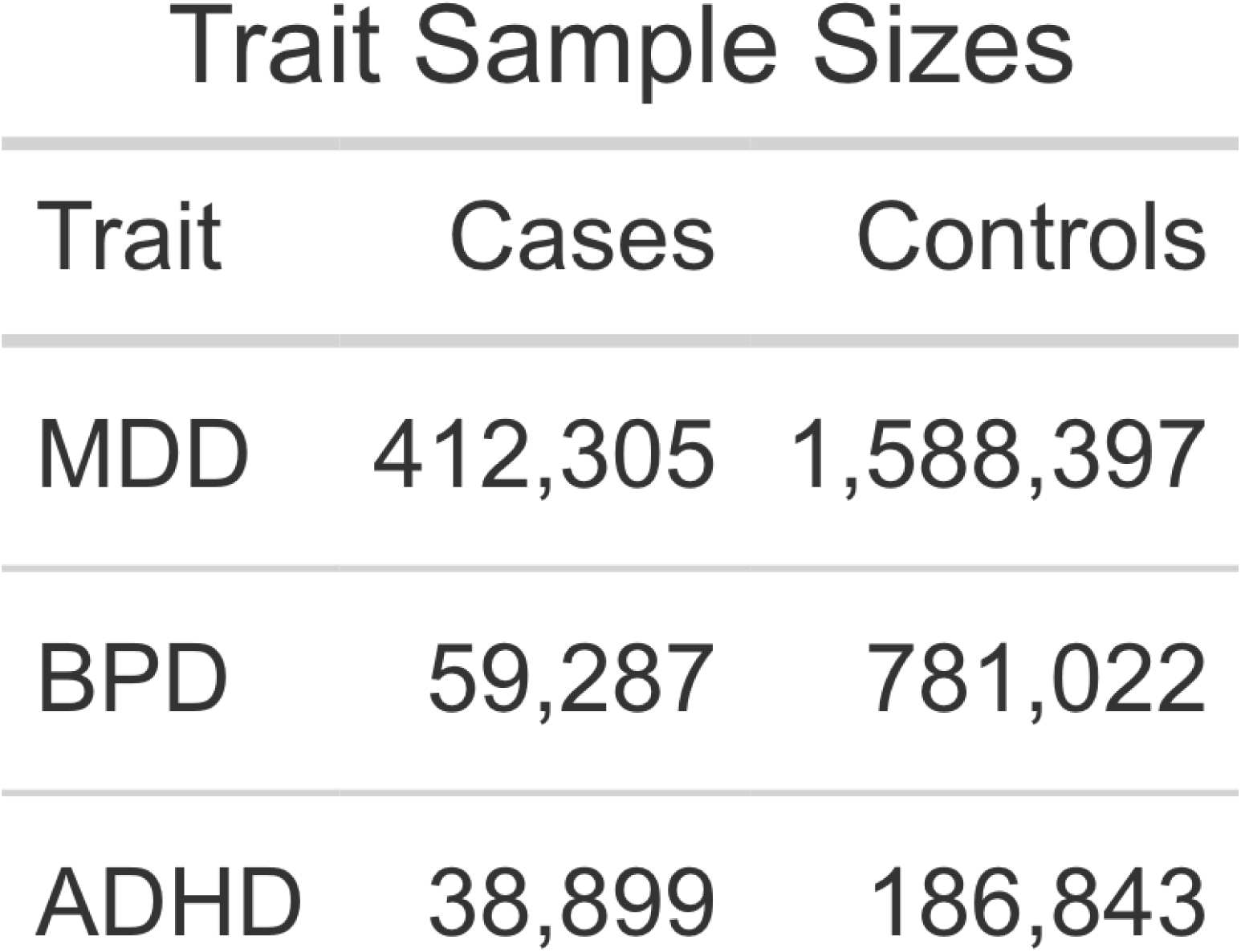
GWAS summary statistics sample sizes from the <u>Psychiatric Genomics Consortium</u>.

**Fig 2.**
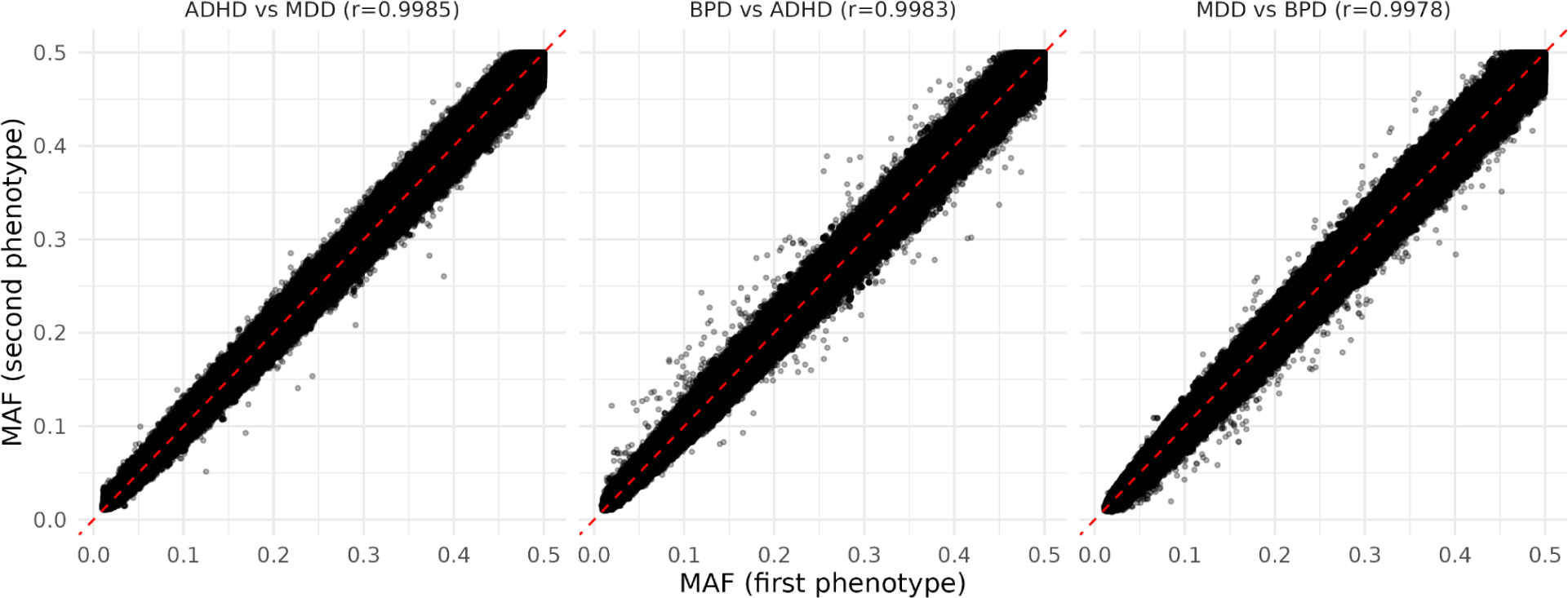
Plot of cohort vs. cohort minor allele frequencies (MAF), with Pearson correlations

**Fig 3.**
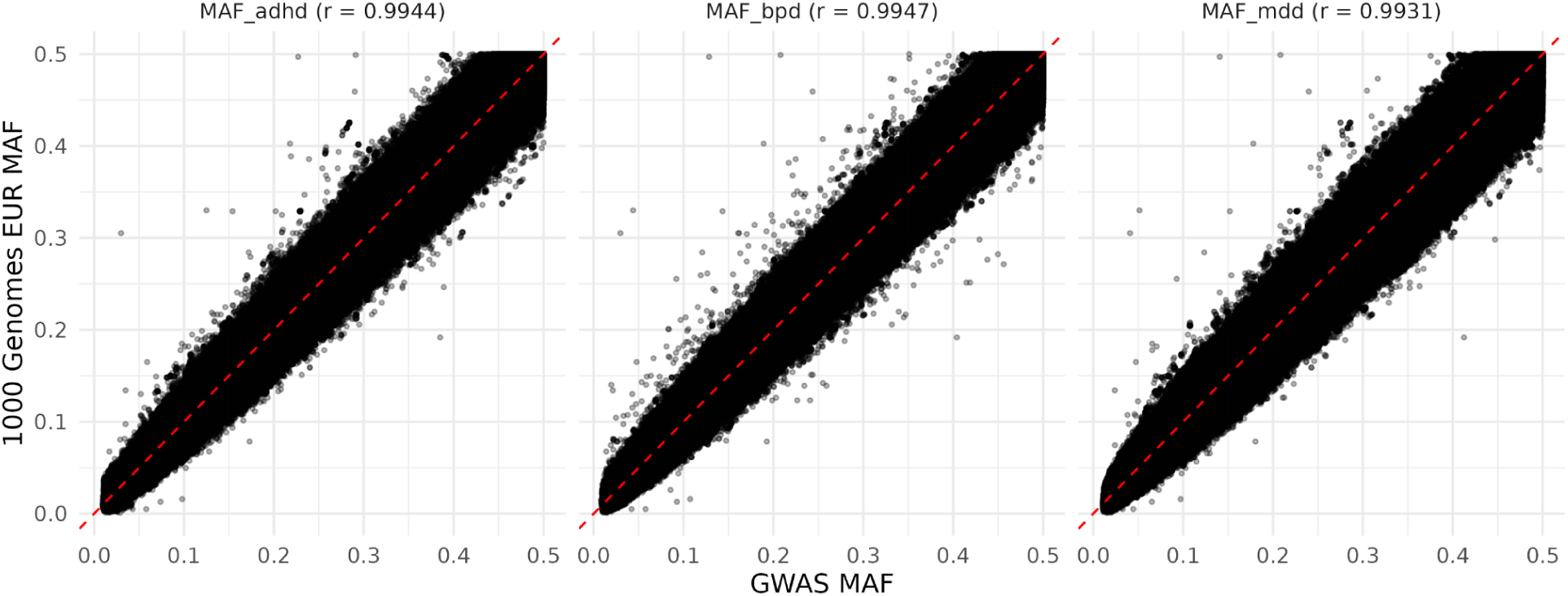
Plot of cohort vs. 1000 Genomes EUR minor allele frequencies (MAF), with Pearson correlations.

### MiXeR

The univariate, bivariate, and trivariate frameworks of MiXeR were used to estimate polygenicity across MDD, BPD, and ADHD. MiXeR is a causal mixture model that assumes a small fraction of variants have an effect on a trait(s), while the remaining variants have zero effect. These causal variant proportions are estimated as parameters by maximizing a log-likelihood function, with the objective of estimating parameters that best describe the observed z-score distributions of input GWAS summary statistics. The model of observed z-score distributions is weighted per-SNP by effective sample size, adjusting for sample size disproportionality. Additionally, the model’s residual error term is specified as a zero-mean, multivariate normal random vector with a freely estimated residual variance-covariance matrix that accounts for sample overlap and cryptic relatedness. The univariate, bivariate and trivariate models of MiXeR work under similar assumptions and allow us to conceptualize the genetic architecture of MDD, BPD, and ADHD by estimating both the number of causal variants needed to explain 90% of SNP heritability and causal variant discoverability (Frei et al. 2019) (Shadrin et al. 2024).

### Genomic Structural Equation Modeling

To gain functional insights into the shared genetic liability across MDD, ADHD, and BPD, the multivariate GWAS and TWAS extensions of genomic structural equation modeling (Genomic SEM) were used. The frameworks utilize structural equation modeling to identify variants and tissue-specific genes that likely underlie the shared covariance across MDD, BPD, and ADHD. This process took place in two steps. The first step estimated total liability scale SNP heritabilities and genetic correlations between each trait using multivariable linkage disequilibrium score regression (LDSC). In the second step, a model is specified such that an unobserved variable F1, drives the shared genetic signal across ADHD, MDD, and BPD. In minimizing a weighted least squares objective, the chosen system of multiple regression and respective covariance associations across the disorders and F1 are estimated as model parameters that best align with the observed covariances from the univariate GWAS summary statistics.

Genomic SEM populates a sampling covariance matrix using a jackknife resampling procedure with sampling errors of total liability scale SNP heritabilities and genetic covariances as diagonals and covariances of sampling errors on off diagonals. The sampling covariance matrix is utilized in a sandwich correction to robustly correct model parameter standard errors, accounting for unknown degrees of sample overlap. Additionally, each trait in the WLS objective is weighted by the inverse diagonals of the genetic sampling covariance matrix, accounting for sample size disproportionality. The trait-specific betas of each individual SNP or gene are then regressed onto the model to estimate an effect size on F1(Grotzinger et al. 2019). Strict quality control procedures were applied to the F1∼SNP summary statistics for downstream analysis, including filtering out SNPs that pass X^2^ and Q_SNP_ genome-wide significance thresholds (p-value < 5e-8). PLINK was used to identify linkage disequilibrium independent SNPs (r^2^<0.1) (Purcell et al. 2007).

### MAGMA / H-MAGMA / MAGMA Cell Typing

Secondary bioinformatics analysis was carried out on the F1∼SNP summary statistics. Hi-C coupled MAGMA (H-MAGMA) was utilized to aggregate F1∼SNP effect sizes to F1∼ gene and gene-set effect sizes (De Leeuw et al. 2015) (Sey et al. 2020). The midbrain, dopaminergic annotation was used to assess the shared genetic liability across MDD, BPD, and ADHD specifically for dopamine neurons (De Leeuw et al. 2015) (Sey et al. 2020). The top 30 protein coding gene hits were subject to further biological analysis.

To examine brain region and cell type associations for F1, MAGMA Cell Typing was used in tandem with the RNA expression dataset curated by Siletti et al. 2023, which consists of over 3 million cell nuclei from 100 locations across the forebrain, midbrain, and hindbrain (Watanabe et al. 2019) (Silletti et al. 2023) (Skene et al. 2016) (De Leeuw et al. 2015) (Murphy, Schilder, & Skene, 2021).

## Results

### Estimating polygenicity with MiXeR

The univariate, bivariate, and trivariate frameworks of MiXeR were used to model shared and trait specific causal variants across MDD, BPD, and ADHD, as well as estimate causal variant discoverability. ADHD’s estimated number of causal variants to explain 90% of SNP heritability (nc@9p) was 7.81K (SD = 291) variants with a causal variant variance (Causal variant discoverability) of 3.00e-05 (SD = 1.03e-06). BPD’s estimated nc@p9 was 7.96K (SD = 254) variants with a casual variant discoverability of 2.64e-05 (SD = 7.35e-07). MDD was the most polygenic and exhibited the smallest causal variant effect sizes across the three disorders, with an nc@9p of 12.18K (SD = 346) variants and a causal variant discoverability of 6.69e-06 (SD = 1.55e-07). The nc@9p for F1 was 11.03K (SD = 190) variants with a discoverability of 1.09e-05 (SD = 1.12e-07).

### Estimating polygenicity with MiXeR - Bivariate

There is an extensive amount of bivariate genetic overlap across the disorders (Fig. 4). MDD exhibited strong genetic correlation with ADHD and BPD. However, despite BPD and ADHD having over 80% nc@9p overlap, a genetic correlation of 0.28 suggests that shared causal variants influence the disorders in ways that are loosely correlated.

**Fig 4.**
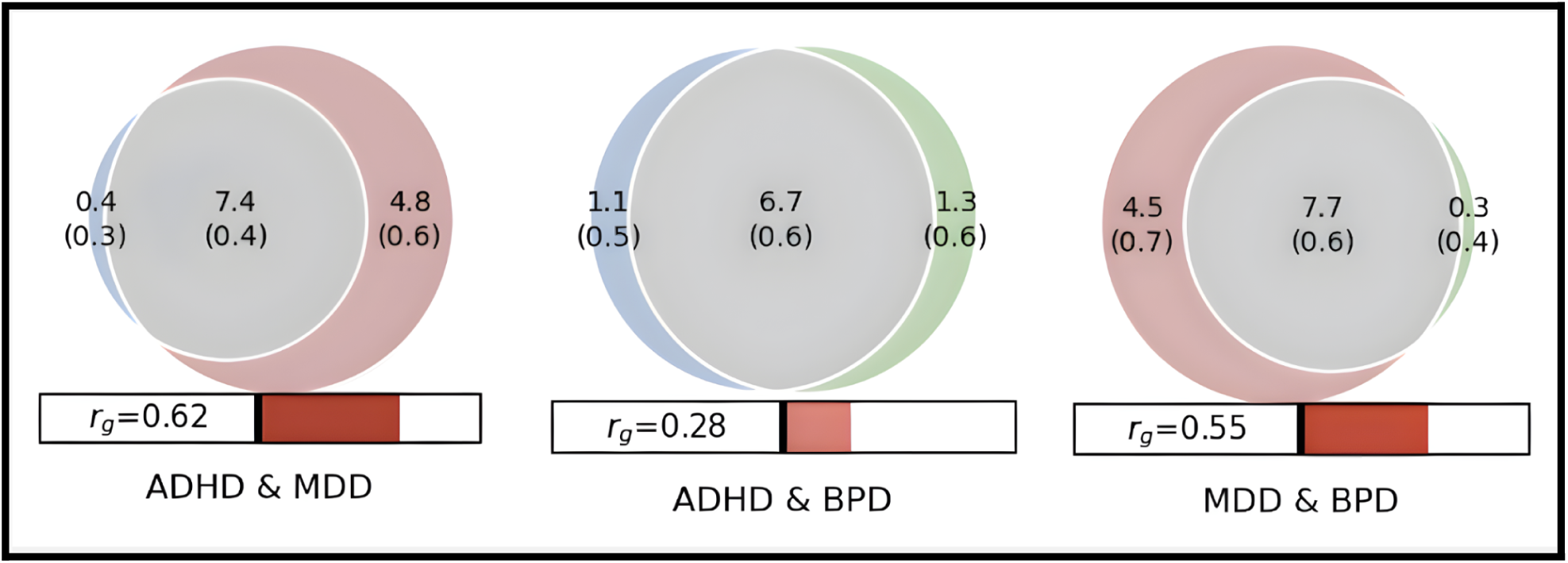
Bivariate genetic overlap across ADHD (Blue), MDD (Red), and BPD (Green), with MiXeR estimates of genetic correlation. Shared genetic variants are colored grey. Venn diagram numbers represent the amount of causal variants (In thousands), that explain 90% of SNP heritability with standard deviations in parenthesis.

### Estimating polygenicity with MiXeR - Trivariate

There is a large degree of a trivariate genetic overlap across MDD, BPD, and ADHD (Fig 5 & 6). The trivariate mixture model estimates highlight ADHD as a potentiator of comorbidity, as consistent with the literature. In the three-way mixture model, less than 1% of the nc@9p variants that make up ADHD are intrinsically specific to ADHD. A majority of the causal variants that influence ADHD are not unique to the disorder itself, but are shared with MDD and BPD.

**Fig 5.**
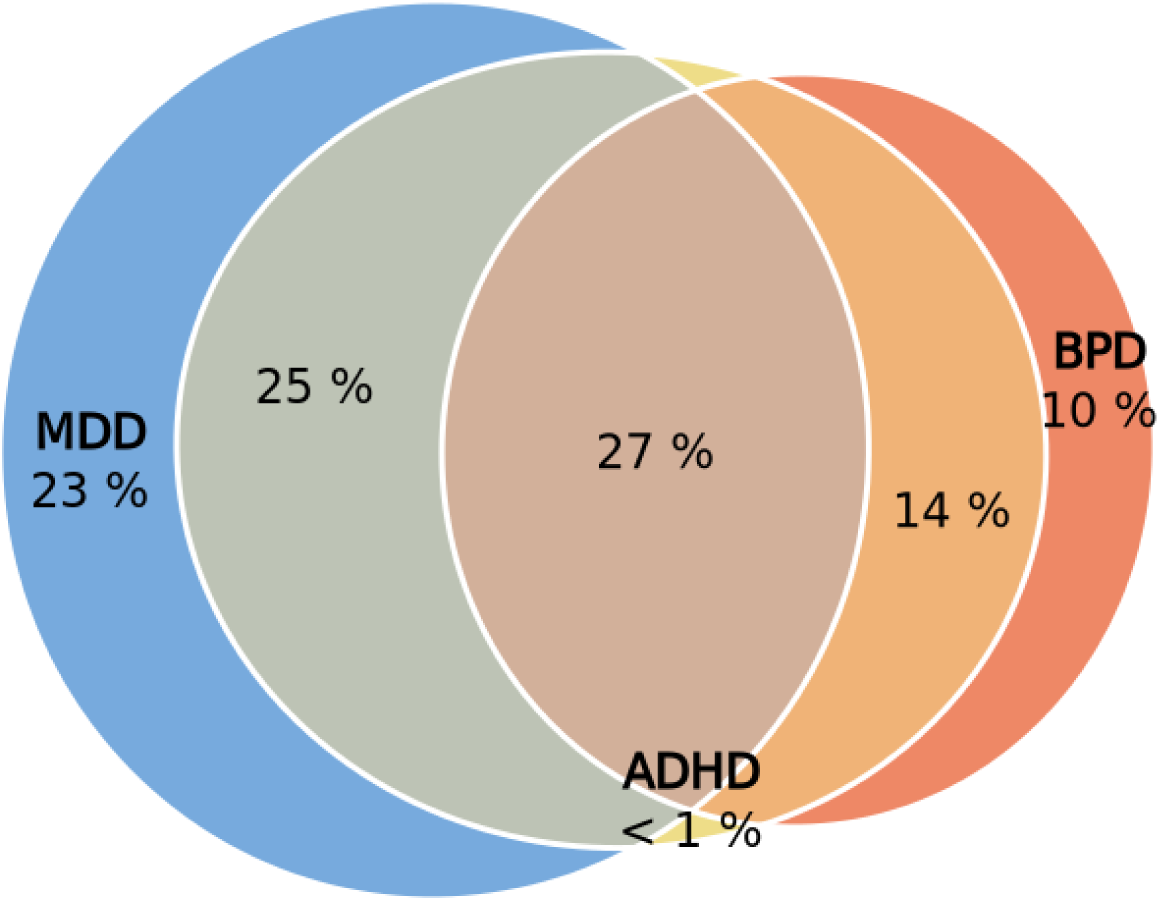
Euler diagram from trivariate mixer analysis of ADHD, BPD, and MDD, denoting the percentage distribution of shared and trait-specific genetic causal variants across the 3 psychiatric disorders.

**Fig 6.**
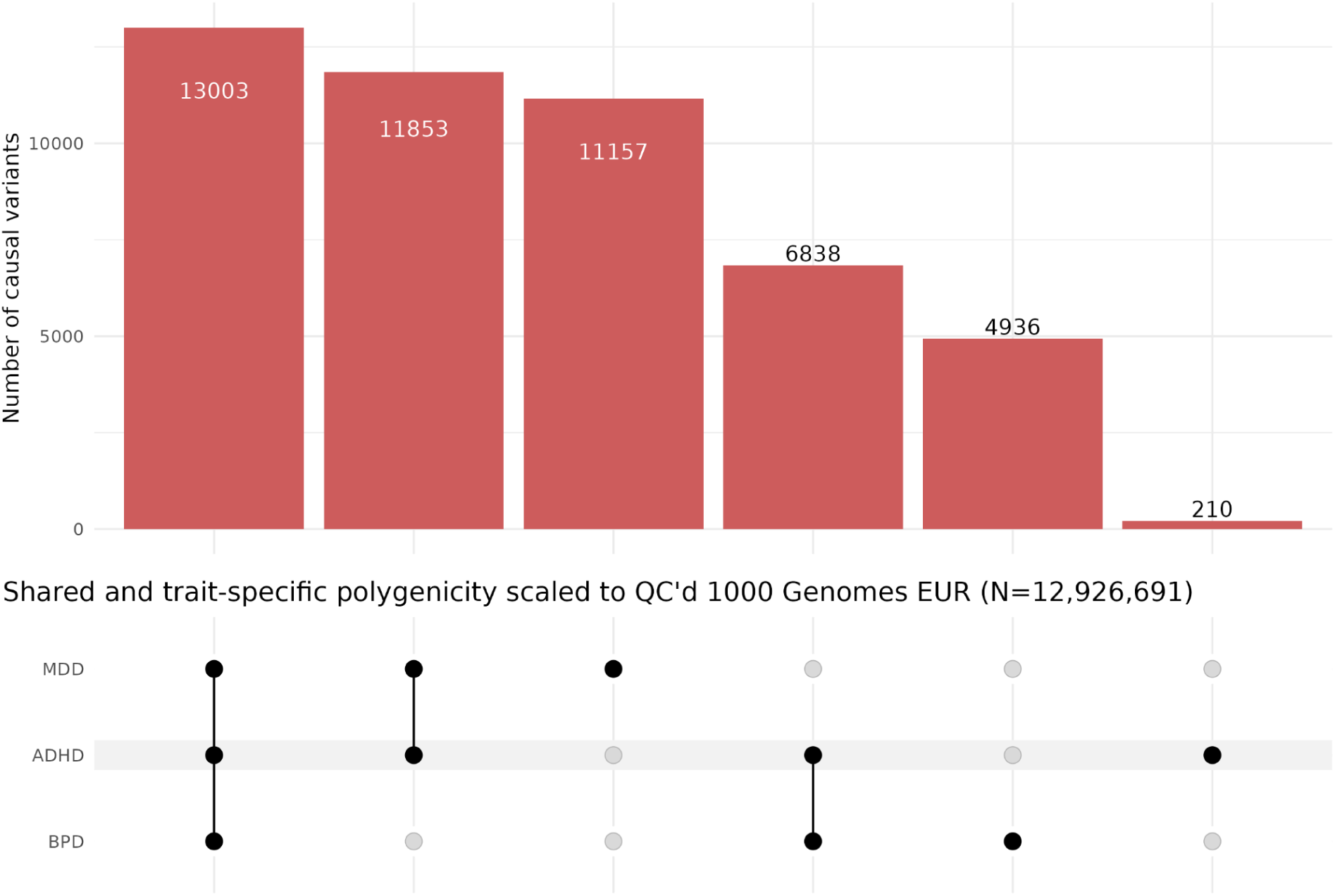
UpSet plot of shared and trait-specific polygenicity across ADHD, MDD, and BPD, scaled to QC’d 1000 Genomes European reference panel (MAF > 0.001, INFO > 0.8, call rate > 0.9, and hardy weinberg equilibrium p-value > 5e-10, N=12926691), from trivariate mixer causal variant proportions.

**Fig 7.**
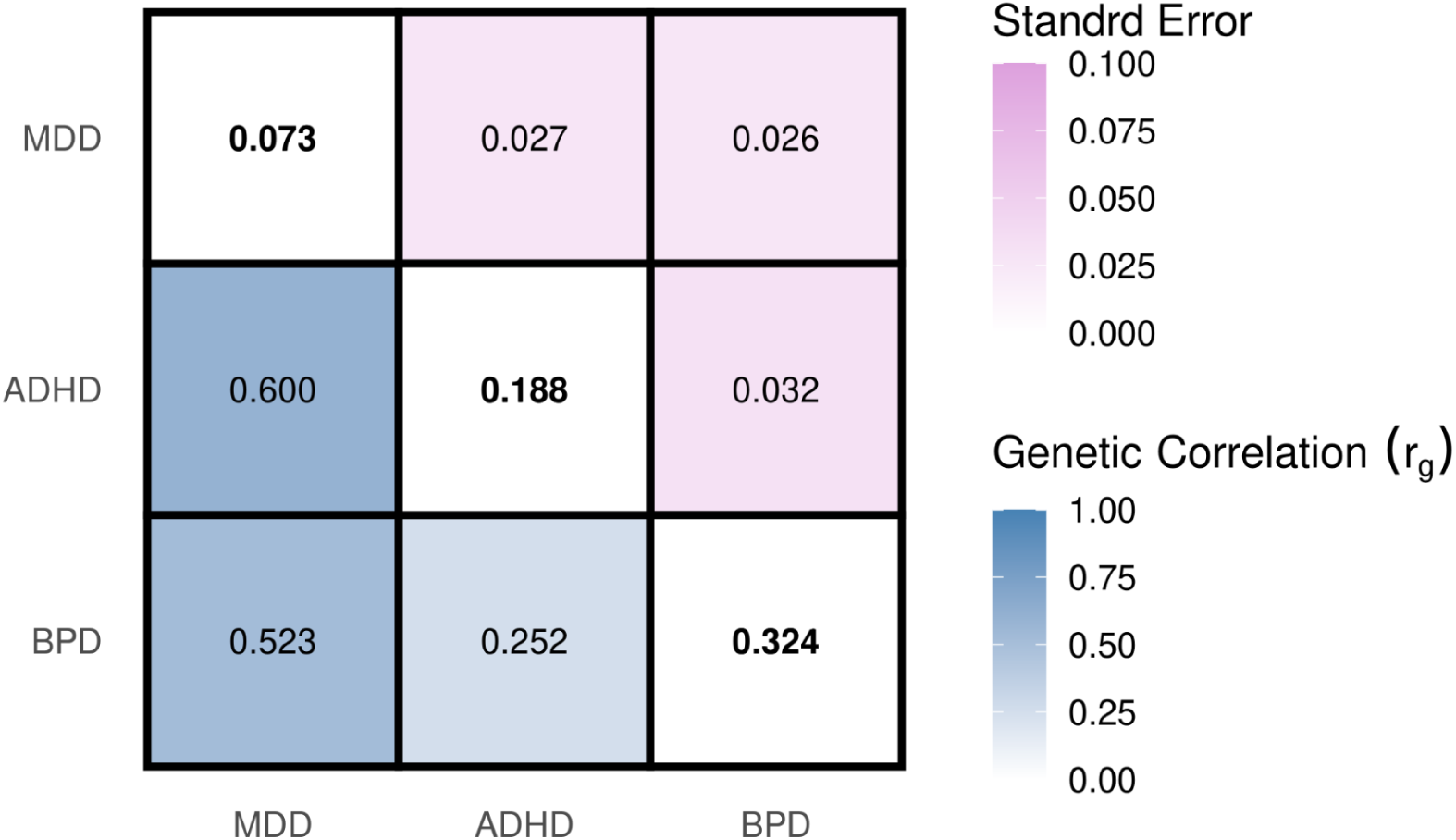
Matrix of total liability scale bivariate genetic correlations (*r_g_*) attained from LDSC in Genomic SEM, where diagonal elements are total liability scale SNP heritability (*h_2_*), bottom off diagonal elements are total liability scale bivariate genetic correlations, and top off diagonal elements are bottom off diagonal standard errors.

**Fig 8.**
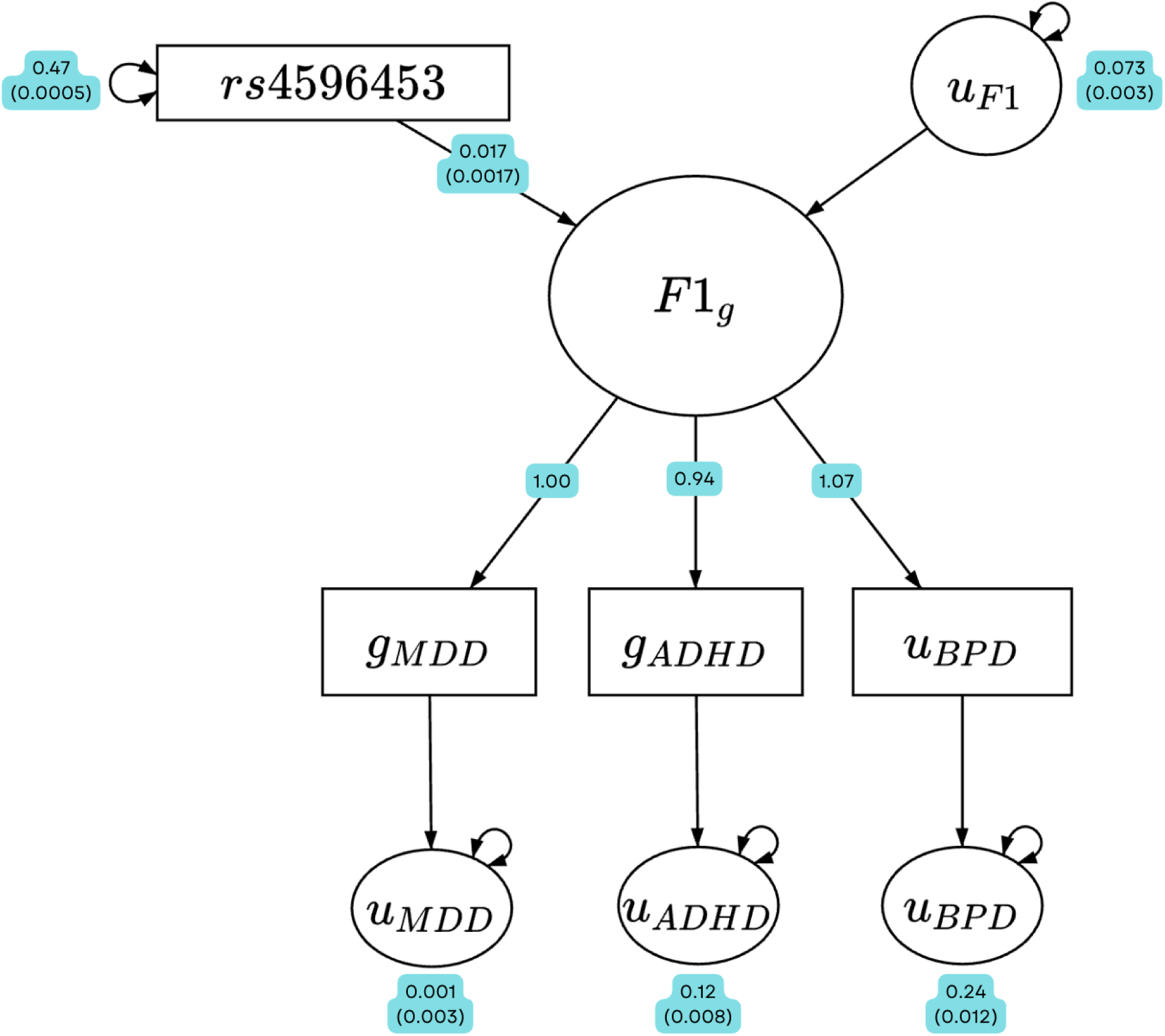
Path diagram depicting how index SNP rs2585813 influences the latent factor F1 from genomic SEM analysis using WLS estimation. All path coefficients are standardized with standard errors in parentheses. In the diagram, “g” subscripts denote genetic variables and “u” subscripts denote the residual variance of their respective genetic variables not explained by F1. The effect of SNP rs2585813 on F1 was 0.017 (SD = 0.00017).

### Gene, gene-set, and celltyping analysis

Utilizing the midbrain dopaminergic Hi-C enriched H-MAGMA annotation, F1 was enriched in 1384 genes (Bonferroni correction, p-value threshold of 9.7e-07) (Fig 9). Across all three disorders, a total of 1025 novel gene associations were revealed in the F1 H-MAGMA analysis that did not reach genome-wide significance in univariate analysis (Fig 15). Using MAGMA’s default gene annotation that maps SNPs to genes via predefined gene coordinates, including SNPs 35kb upstream and 10kb downstream, a total of 28 gene sets reached genome-wide significance (Fig 10). A significant proportion of genome-wide significant gene-sets are involved in various dynamics of the neuronal synapse. In all brain regions and cell-types assayed, F1 was enriched to varying degrees. Most notably, F1 was significantly enriched in the cerebellum and in medial ganglionic eminence interneurons. In the multivariate transcriptome-wide analysis, F1 was enriched in a total of 272 genes across GTExv8 brain regions (P-value < 1e-06) (Fig 14).

**Fig 9.**
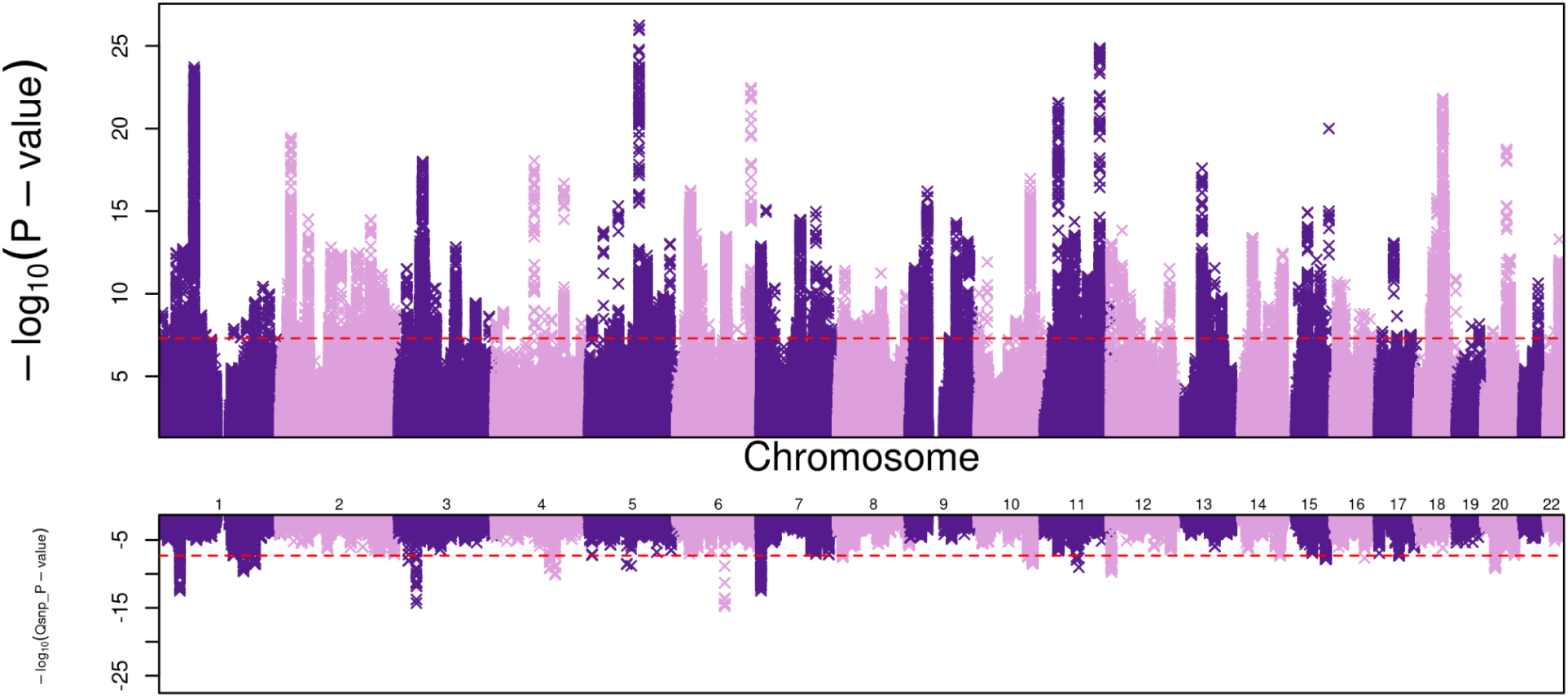
Miami plot of genome wide association results for F1 with p-value thresholds of 5e-08 for both panels, represented by the red dotted line. The top panel displays the -log_10_(p-value) for each SNP, and the bottom panel displays the SNP’s respective -log_10_(Q_SNP_ p-value) from the heterogeneity test (Q_SNP_ test estimates the degree to which a SNP is not mediated by F1).

**Fig 10.**
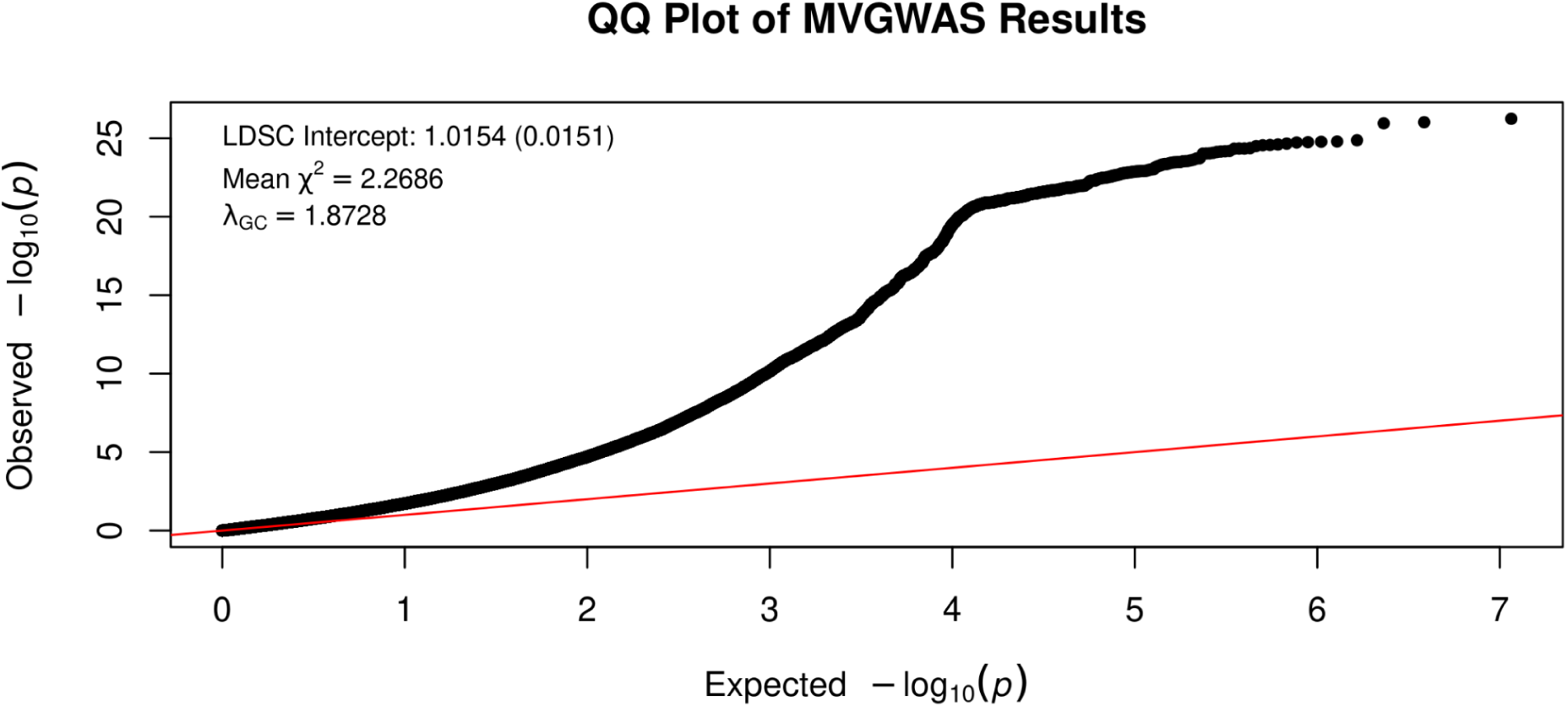
QQ plot of expected vs. observed -log_10_(p-value) for the F1 summary statistics with LDSC intercept, mean chi-squared, and lambda.

**Fig 11.**
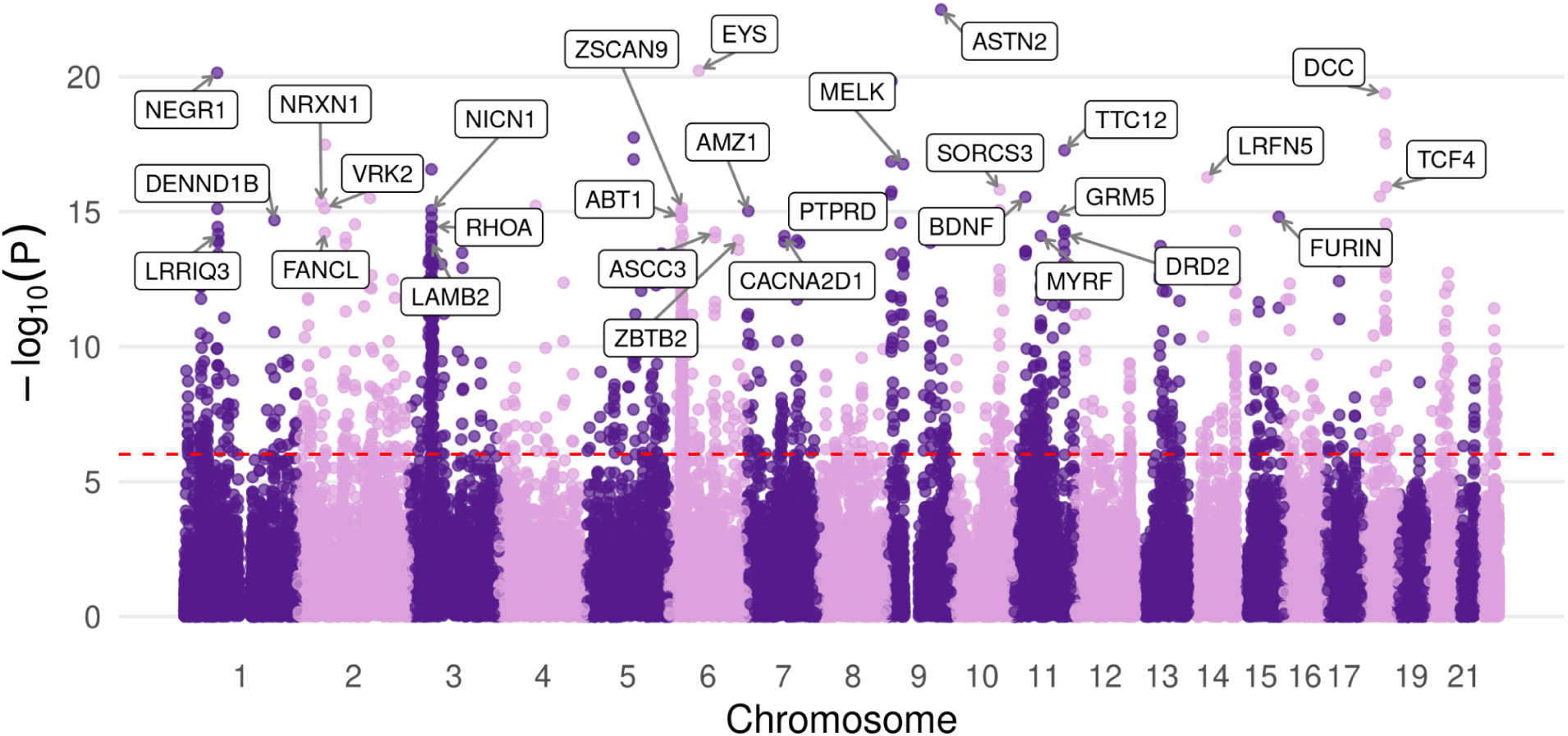
Manhattan plot of F1 gene associations from DA neuron H-MAGMA analysis. The top 30 protein coding genes are labeled, and the Bonferroni adjusted p-value threshold is represented by the dotted line (p=9.7e-07).

**Fig 12.**
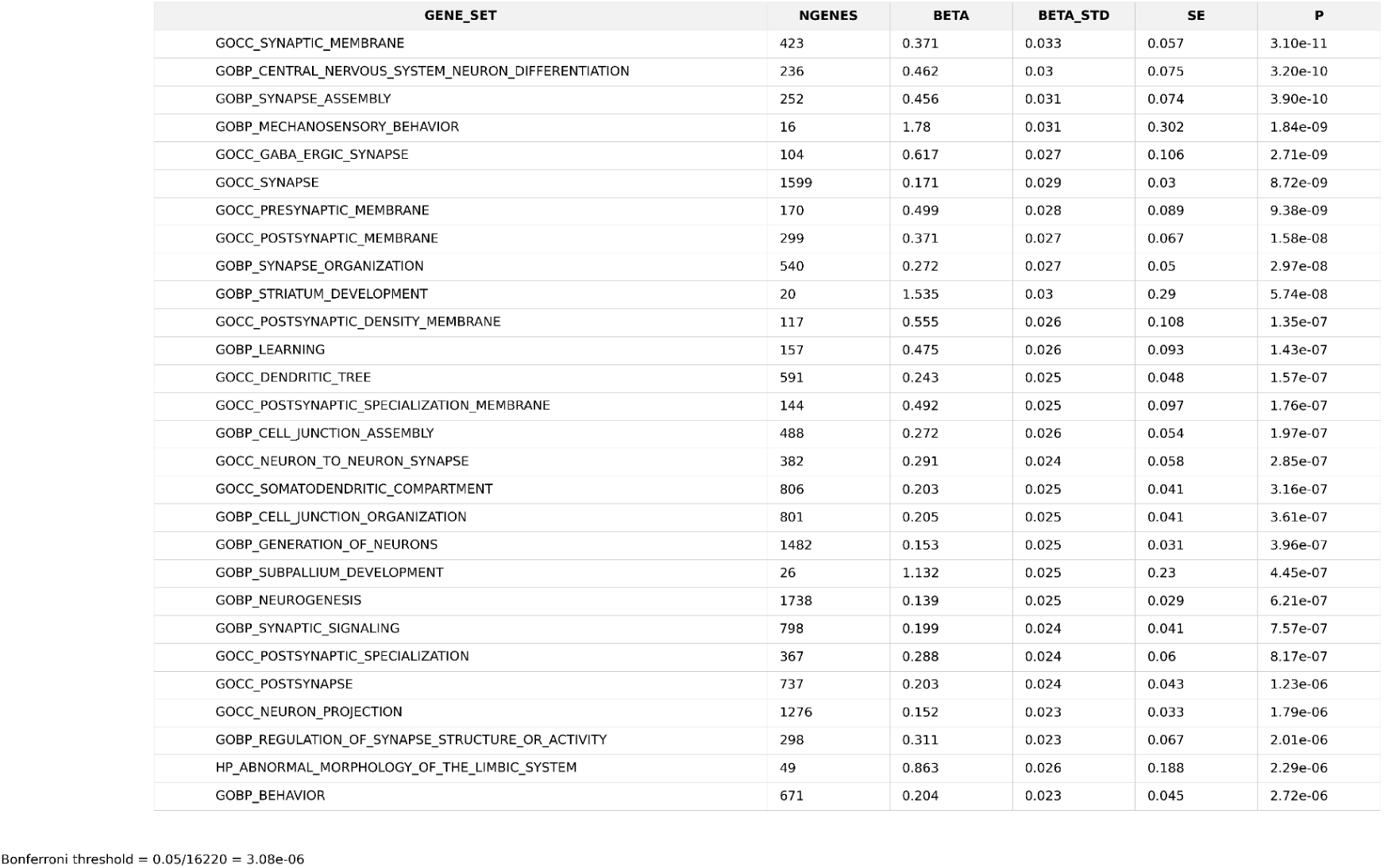
Table of F1 genome-wide significant gene-sets using MAGMA’s default gene annotation file.

**Fig 13.**
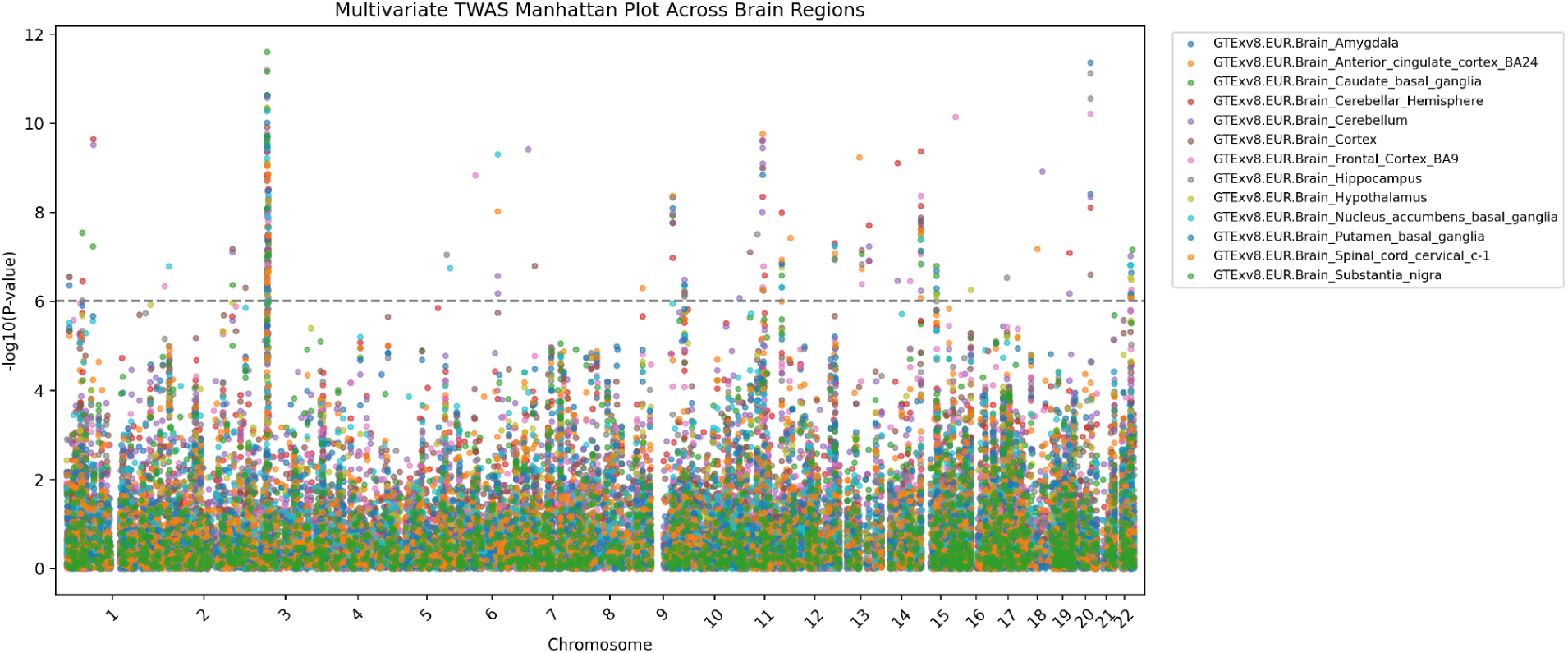
Manhattan plot of F1 TWAS gene associations from GTExv8 brain regions. The significance threshold is denoted by the grey dotted line.

**Fig 14.**
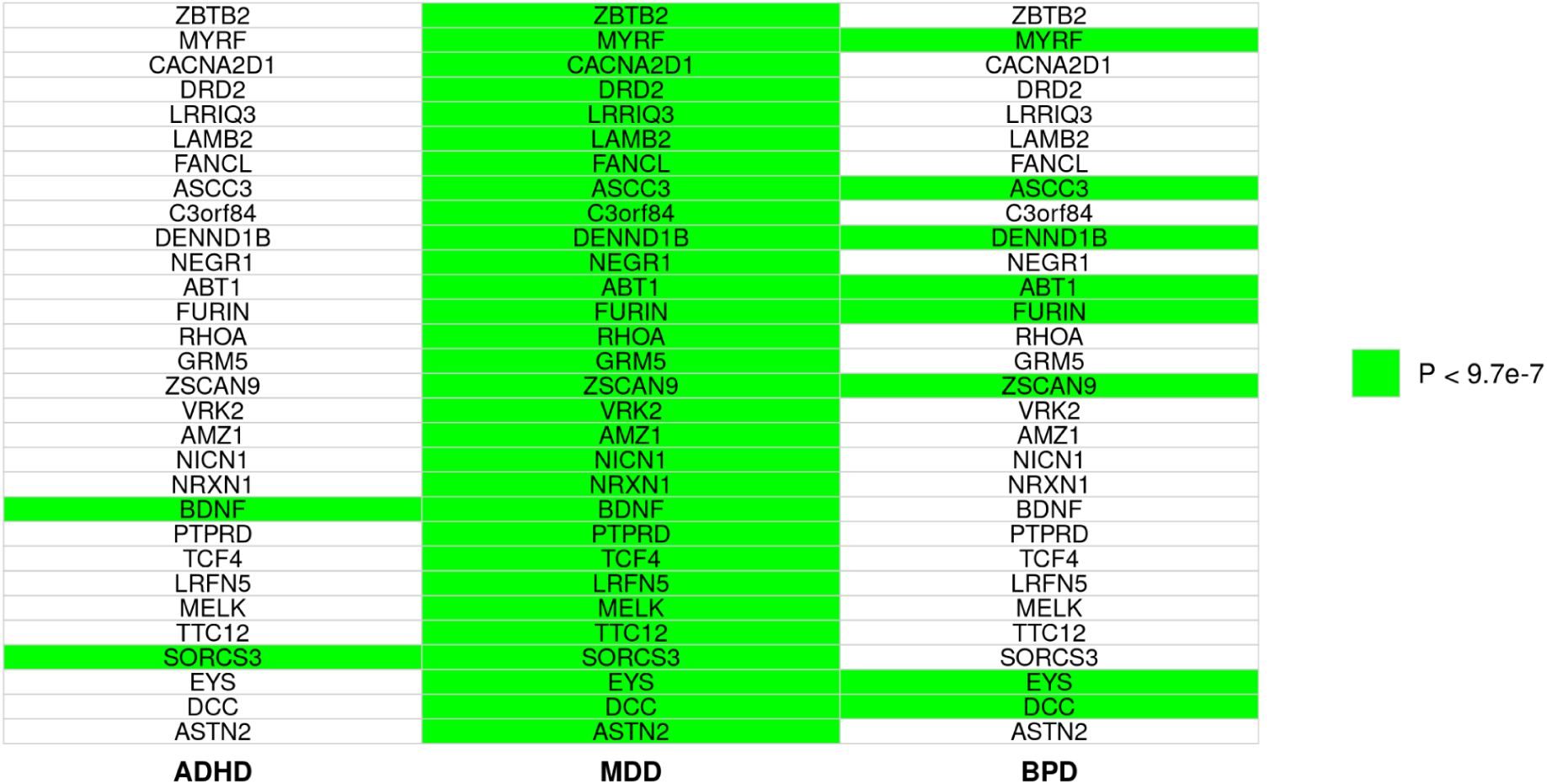
List of the top 30 protein-coding genes associated with F1 from the H-MAGMA gene analysis, showing for each gene whether it reached genome-wide significance in the univariate GWAS of ADHD, MDD, and BPD. Genes highlighted in green denote p-values exceeding significance thresholds for the respective trait.

**Fig 15.**
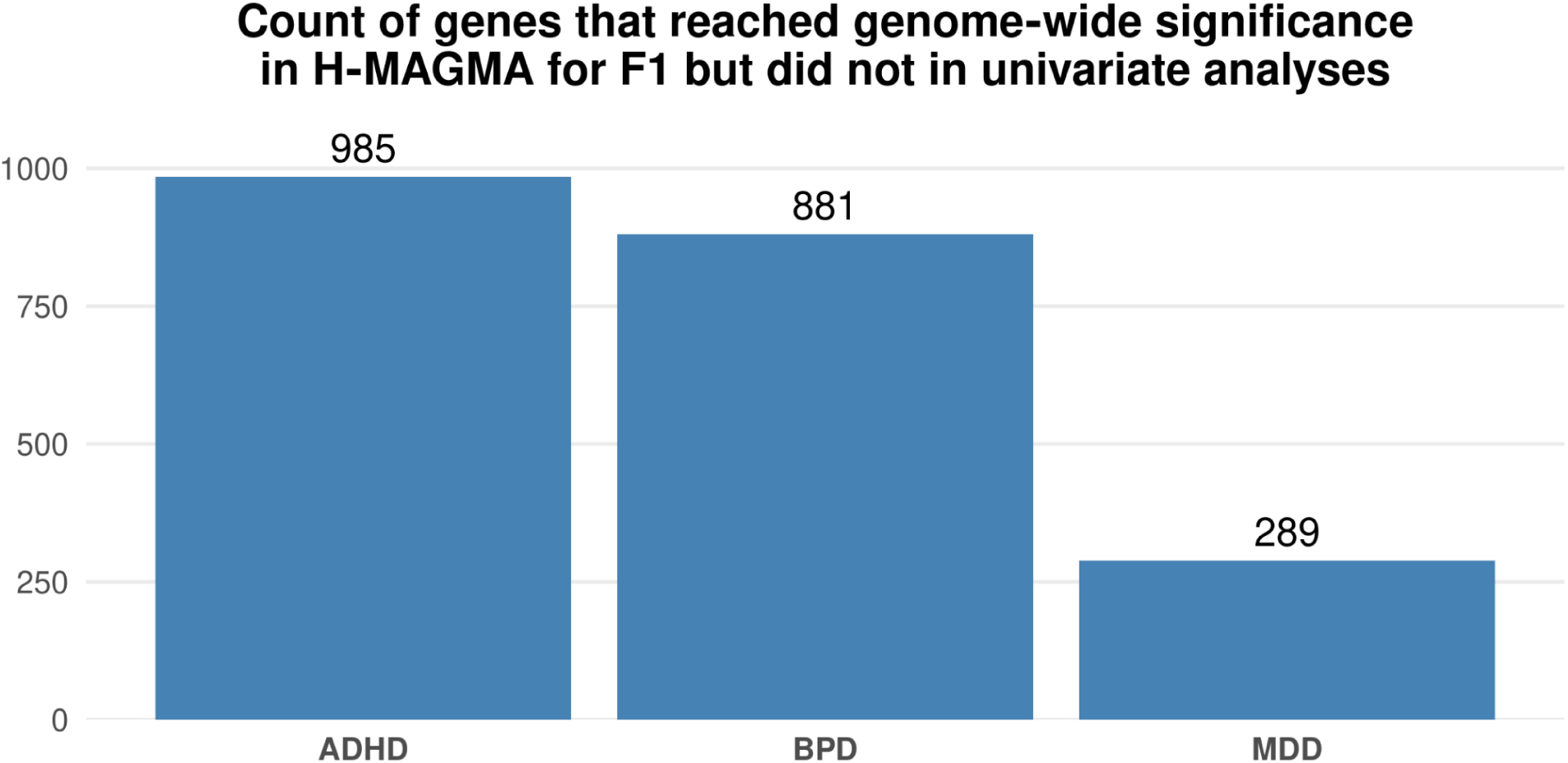
Bar chart summarizing F1 gene discovery that went undetected in univariate DA neuron H-MAGMA analysis. The chart depicts the number of genes that reached genome-wide significance in the dopaminergic H-MAGMA analysis for F1 but not for the univariate summary statistics of BPD, ADHD, and BPD.

**Fig 16.**
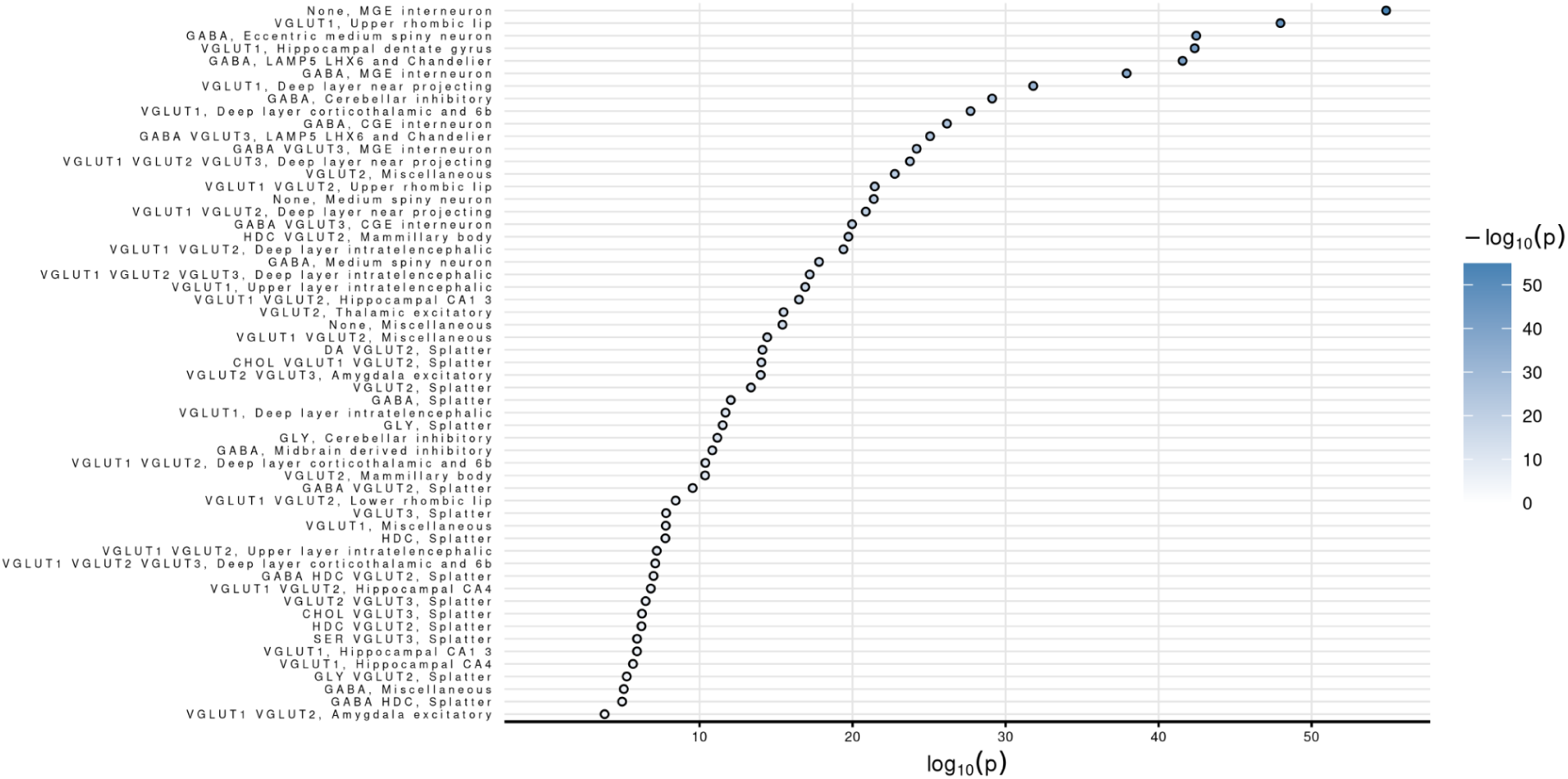
Dotplot of cell types associated with F1 from MAGMA Cell Typing analysis using the Human Brain Atlas v1.0 RNA expression dataset.

**Fig 17.**
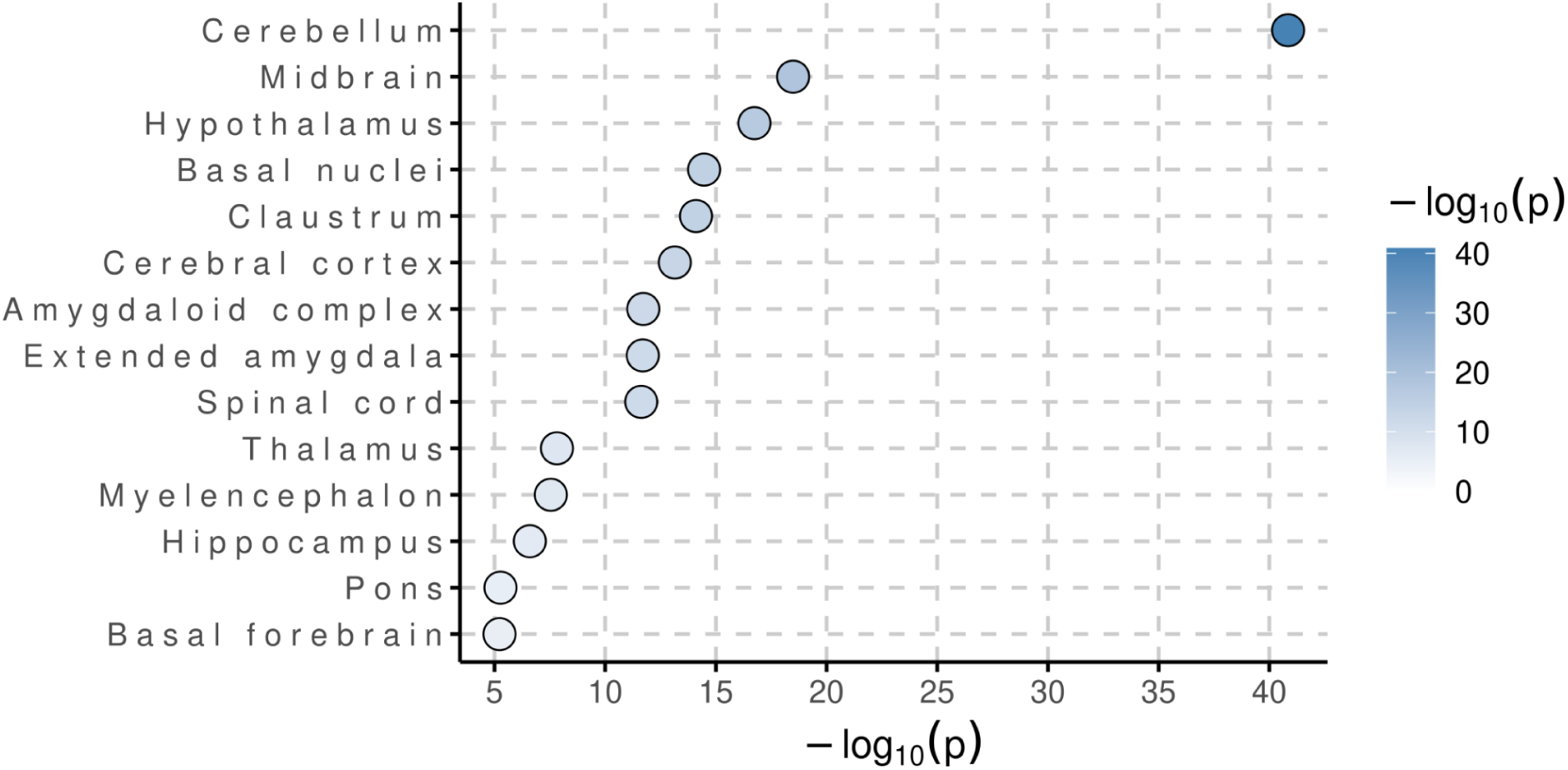
Dotplot of brain regions associated with F1 from MAGMA Cell Typing analysis using the Human Brain Atlas v1.0 RNA expression dataset.

## Discussion

In the context of multivariate GWAS-driven genetic discovery, there remains a substantial amount of undiscovered associations (Turchin & Stephens, 2019). This is especially true for GWAS that have small sample sizes, for which a great number of causal SNPs do not reach genome-wide significance due to lack of power. However, we have demonstrated that multivariate frameworks like genomic SEM can expedite genetic discovery for underpowered GWAS studies by leveraging genetic covariances. Over 80% of the case sample size of the multivariate GWAS originates from the MDD summary statistics. We were able to leverage the effect size of MDD to reveal 1025 DA neuron gene associations for BPD and ADHD that went undetected in univariate analyses.

### Candidate gene biological annotation - *PTPRD*

We chose to conduct an in depth review of one of the top DA neuron H-MAGMA hits, evaluating its biological plausibility as a risk gene for F1. Protein tyrosine phosphatase receptor delta (*PTPRD)*, was among the top 10 protein coding genes associated with F1, p = 2e-16 (0.05/51684 genes tested = p-value threshold of 9.67e-07). Given *PTPRD*’s cell adhesion mediating immunoglobulin and fibronectin III domains, and its catalytically active D1 phosphatase domain, we identified *PTPRD* as a possible risk gene (Uhl & Martinez, 2019).

Given that neurotransmission homeostasis is heavily implicated in neurological disorders, we hypothesize that *PTPRD* may regulate DA neurotransmission, from which dysregulation and or dysfunction contribute to the development of MDD, ADHD, and BPD (Ioana Teleanu et al. 2022).

### Review of *PTPRD* KO mice studies and DA-related behavioral alteration

Among all studies that assay reward-related behaviors, *PTPRD* KO/HET/Inhibited mice exhibited significant deviation from the wild-type (Uhl et al. 2018) (Drgonova et al. 2015) (Ho et al. 2023) (Uhl & Martinez, 2019). We then asked how and why *PTPRD* knockout and inhibition significantly altered reward-related behaviors such as cocaine-conditioned place preference, cocaine self-administration, motivation for cocaine, and goal oriented behavior particularly in the context of DA’s involvement in reward processing and *PTPRD*’s association with F1 in the dopaminergic Hi-C enriched gene analysis.

### PTPRD/TrkB and PDGFRB

PDGFRB and TrkB are well documented substrates of *PTPRD*’s D1 phosphatase domain (Uhl & Martinez, 2019) (Xu et al. 2022). In the absence of *PTPRD*, *TrkB* and *PDGFRB*, which potentiate downstream the *MEK*-*ERK* pathway, remain hyperactivated. Higher concentrations of downstream effectors, *MEK* and *ERK*, are observed in *PTPRD* KO/HET mice when compared to the WT (Toita et al. 2020). Due to *PDGFRB*’s negligible expression in human dopaminergic RNA expression datasets, we excluded the gene from our biological annotation (Supplementary Note 3 & 4).

### ERK1/2 signaling involved in regulation KCNQ2 K+ ion channels

*ERK1/2* is heavily implicated in the LTP of neurons in the nucleus accumbens in response to dopaminergic signaling (Nagia et al. 2016). Dopamine-mediated signaling of the D1 receptor activates the *PKA*/*RAP1*/*ERK* and inhibits KCNQ-mediated K+ current via direct phosphorylation of *KCNQ2* ion channels (Nagia et al. 2016) (Tsuboi et al. 2022). Elevated ERK1/2 signaling in the absence of *PTPRD* may alter the ability of DA neurons to maintain excitability homeostasis via *D1R*/*ERK*/*KCNQ2* signaling (Supplementary Note 5).

### *BDNF*/*NTRK2*’s role in regulating survival and neuroplasticity

*BDNF*, *TrkB*’s canonical ligand, is a promoter of DA neuron survival and axonal growth, regulating survival and neuroplasticity of DA neurons (Østergaard et al. 1996) (Hydman et al. 1991). Dopamine neurons require *BDNF* to undergo LTP from glutamatergic excitatory synapse input (Pu et al. 2006). Upon *BDNF* stimulation, decreased *Rac1*-GTP concentrations are observed in *VAV2/3* -/- mice when compared to the wild type, suggesting that *VAV2*’s GEF domain is required for *Rac1*-GTP synthesis. Additionally, post administration of theta-burst stimulation at CA1 glutamatergic synapses, significantly reduced long term potentiation was observed in *VAV2/3* deficient mice when compared to wildtype (Hale et al. 2011). In the absence of PTPRD, hyperactivation of *TrkB*/*VAV2* signaling may result in excessive dendritic spine head growth, leading to chronic excitability of DA neurons (Supplementary Note 6-8)

### DA neuron BDNF/TrkB signaling and inhibitory LTD of the GABAergic synapse

BDNF acts via TrkB to potentiate retrograde, endocannabinoid-mediated inhibitory LTD upon otherwise subthreshold stimulation from GABAergic synaptic currents (Zhong et al. 2015). The absence of *PTPRD* may result in excessive inhibition of GABAergic synapses via hyperactivated *TrkB* signaling, contributing to the chronic excitation of DA neurons (Supplementary Note 9).

### PTPRD deficient mice experience enhanced long-term potentiation

A common trend we have identified from the potential downstream effectors of *PTPRD* is chronic excitability. These findings are consistent with *PTPRD* KO/Inhibition literature that assay neuronal excitability, of which there is a singular publication. As the current literature suggests, core pathways like *TrkB*/*BDNF* and *MEK*/*ERK* are typically conserved across neurons (Gupta et al., 2013) (Hirata et al., 2017). It is biologically plausible that *PTPRD* may play non-redundant roles in DA neuron LTP and inhibitory LDP, from which strict homeostatic control is required to achieve optimal reward processing (Supplementary Note 10). When neurons are chronically excited, the signal threshold for inducing LTP increases, a phenomenon known as metaplasticity. The absence of *PTPRD* and hyperactivation of *TrkB* signaling may result in chronic excitation, increasing LTP and I-LDP signaling thresholds and attenuating cocaine’s ability to induce both LTP and I-LDP in dopamine neurons and ultimately reinforce reward-related behavior (Supplementary Note 11).

### VAV2 and DAT transporter trafficking in the mesolimbic pathway

LAR-family RPTPs dephosphorylate tyrosine residues involved in RET activation, as well as prevent dimerization to ultimately hinder autophosphorylation. These interactions are independent of GDNF, RET’s canonical ligand, and serve to downregulate RET signaling pathways (Uetani et al., 2009; Qiao et al., 2001). The loss of PTPRD may result in hyperactivated RET signaling. *GDNF* stimulation induces Tyr-905 phosphorylation of RET, which is accompanied by a rise in *VAV2* tyrosine phosphorylation (Zhu et al. 2015). *VAV2* is an essential mediator of DAT transporter membrane trafficking for DA neurons in the ventral tegmental area (Zhu et al. 2015). *TrkB* hyperexcitability and consequent increases in *RET* and *VAV2* tyrosine phosphorylation in the absence of *PTPRD* may result in increased *VAV2*/*RET* mediated endocytosis of *DAT*, which may underlie the observed behavioral responses to cocaine in PTPRD KO/inhibition mice (Supplementary Note 12).

### ERK1/2 and DAT transporter

*ERK1/2* regulates dopamine transporter surface expression and transport capacity (Moron et al. 2003). In the context of *PTPRD* KO/inhibition, it is important to note that the literature is mixed in supporting elevated phosphorylation of *ERK1/2*. Conditional mice knockouts of *PTPRD* in *EMX1*+ neural progenitor cells only increased phosphorylation of *MEK1/2*, whereas global mice knockouts of *PTPRD* increased both phosphorylation of *MEK1/2* and *ERK1/2* (Cortés et al. 2024). However, it is plausible that *PTPRD* inhibition in DA neurons hyperactivates *ERK1/2* signaling, resulting in chronic excitability from excessive deactivation of *KCNQ2* ion channels and aberrant homeostatic control of *ERK1/2*-mediated surface *DAT* expression and phosphorylation (Supplementary Note 13).

### PKC signaling and DAT

PKC is a downstream effector of *BDNF* signaling and heavily implicated in DAT transporter trafficking (Gabriel et al. 2013) (Lanuza et al. 2019). The absence of *PTPRD* may result in hyperactivation of PKC signaling, attenuating the ability of DA neurons to maintain homeostatic control of DAT surface levels (Supplementary Note 14).

### Lower mRNA and protein concentrations of PTPRD concomitantly accompanied by lowered mRNA and protein concentrations of DAT transporter in RLS mice model

There is extremely limited research on *PTPRD*’s involvement with *DAT* transporter regulation, however in an animal model of restless leg syndrome, spontaneously hypertensive rats exhibited lower mRNA and protein concentrations of *PTPRD*, which was concomitantly accompanied by lower mRNA and protein concentrations of the *DAT* transporter when compared to controls (Morais et al. 2023). The pathogenesis of RLS is highly characterized by dopamine dysfunction and reduced iron concentrations in the brain (Guo et al. 2017). *DAT*-KD mice experience increased synaptic DA and permanent activation of postsynaptic D1 and D2 receptors, ultimately leading to desensitization (Savchenko et al. 2023). Lower concentrations of *PTPRD* may contribute to lower concentrations of the *DAT* transporter, increasing DA under the curve. Sustained increases in extracellular DA concentrations may result in the desensitization of D1 and D2 receptors in the absence of *PTPRD*’s D1 phosphatase domain.

### PTPRD and regulation of hedonia

Anhedonia is heavily implicated in major depressive disorder and in the depressive episodes associated with bipolar disorder (Whitton & Pizzagalli, 2022). Anhedonia is also implicated in ADHD (Sternat et al. 2018). Recent literature consistently observes reduced activation of the ventral striatum during reward processing in people diagnosed with ADHD, BPD, and MDD (Scheres et al. 2007) (Yip et al. 2014) (Pizzagalli et al. 2009). The ventral striatum is heavily implicated in reward processing, and the area is routinely activated during stages of reward processing from ventral tegmental area projections (Liu et al. 2010) (Qi et al. 2016). We hypothesize that PTPRD may be essential for DA neurons to maintain homeostatic neurotransmission in reward-associated circuits to regulate hedonic behavioral response. This hypothesis may underlie alterations to reward-associated behaviors when *PTPRD* and or its downstream effectors are experimentally attenuated. It is plausible that the observed effect size of *PTPRD* for F1 in the DA neuron H-MAGMA analysis has a direct relationship with the reduction of ventral striatum activation as it relates to reward processing in those with MDD, BPD, and or ADHD.

### PTPRD-F1 model proposal

Loss of PTPRD’s D1 phosphatase domain may permit the hyperactivation of *TrkB* and RET signaling pathways, resulting in chronic excitation and aberrant, *DAT*-mediated uptake of synaptic dopamine. Hyperactivated *TrkB* and *RET* signaling may increase *ERK1/2*, *PKC*, and *VAV2* phosphorylation and activity. The elevated occurrence of downstream *VAV2* mediated dendritic spine head growth likely attenuates homeostatic control of structural LTP that respond to glutamatergic input. The elevated downstream occurrence of phosphorylated *ERK1/2* may attenuate homeostatic control of both *DAT* transporter surface expression and *KCNQ2* K+ ion channels. The elevated occurrence of downstream *PKC* and *VAV2* activation may attenuate homeostatic control of bidirectional *DAT* transporter surface expression. In addition, excessive TrkB signaling may induce elevated retrograde inhibition of inhibitory GABAergic synapses.

The coalescence of these *PTPRD* risk factors may lead to the chronic excitation of DA neurons through increases of structural LTP and inhibitory LDP signaling thresholds and decreased DAT-mediated synaptic dopamine uptake, from which the D1 and D2 receptors become desensitized. We propose that these mechanisms underlie the consistent trend of altered behavioral responses to cocaine in *PTPRD*-deficient mice, especially in regards to decreased motivation for cocaine (Supplementary Note 15).

### Exploratory, hypothesis-generating PTPRD fine-mapping using 1000 Genomes external reference panel and F1 summary statistics

The H-MAGMA DA neuron variant annotation mapped SNPs to *PTPRD* from 7.4 Mb to 12 Mb on chromosome 9, suggestive of cis-regulatory elements. We then estimated causal SNP posterior inclusion probabilities and local heritability for F1 in regions of interest, as well as analyzed the LD structure of *PTPRD*. There is a significant downstream spike in LD from about 10.9Mb to 12Mb on chromosome 9 (Supplementary Fig 13). Functionally relevant domains such as expression quantitative trait loci (eQTLs), methylation quantitative trait loci (meQTLs), and chromatin loops exhibit reduced recombination (Liu et al., 2017). Concurrently, there is an F1 risk locus within this elevated region of LD. Using LAVA, we estimated local genetic heritability using the F1 summary statistics as input (Werme et. al 2022) (Supplementary Note 16). We then utilized mixer-finemap to estimate causal posterior inclusion probabilities for each SNP in the risk locus (Akdeniz et al. 2024) (Supplementary Fig 17). Given the concurrence of abnormally high SNP heritability and reduced recombination, this region may be a DA neuron regulatory domain, housing cis-eQTLs for PTPRD and a plethora of other genes (Supplementary 16).

## Acknowledgements

Anna K. Lack^1^ - Research Mentor

## Supplementary Information

Supplementary Information and Supplementary Figures 1-17.

## Data and code accessibility

All scripts used in this paper can be found at https://github.com/chrislaw133/PTPRD. Data generated in this study can be found at https://zenodo.org/records/16741578. All data analyzed in this study can be found in the published articles’s data repositories (See references).

